# Osteoimmunomodulatory Stem Cell Nanoghosts as a Novel Nanotherapeutic for Bone Regeneration

**DOI:** 10.64898/2026.05.11.724218

**Authors:** Giovanni A. Micheli, Tianyu Yang, Debby Gawlitta, Kenny Man

## Abstract

Critical-sized bone defects and implant-associated complications are often exacerbated by chronic inflammation, which compromises tissue repair and implant integration. Mesenchymal stromal cell (MSC)-derived extracellular vesicles have emerged as promising immunomodulatory nanotherapeutics; however, their clinical translation remains constrained by low yield, heterogeneity, and poor scalability. Here we present a bioengineered MSC-derived nanoghosts platform designed to overcome these translational barriers while enabling tunable osteoimmunomodulatory function. By coupling high-yield nanoghost fabrication with biomimetic MSC conditioning, we demonstrate that oxygen tension (5 or 21% O_2_) and 3D culture substrates (5 or 15 wt-% GelMA) can reprogram MSC immunophenotype. Nanoghosts generated under hypoxic and 3D conditions displayed enriched anti-inflammatory cargo, preserved MSC viability under inflammatory stress, and partially rescued osteogenic mineralization in the presence of pro-inflammatory cytokines. Together, these findings showcase MSC nanoghosts as scalable and bioactive immunoregulatory nanotherapeutic capable of modulating immune-bone crosstalk, providing a translational strategy to mitigate inflammation-driven impairment of bone regeneration and implant integration.

**Graphical abstract:** 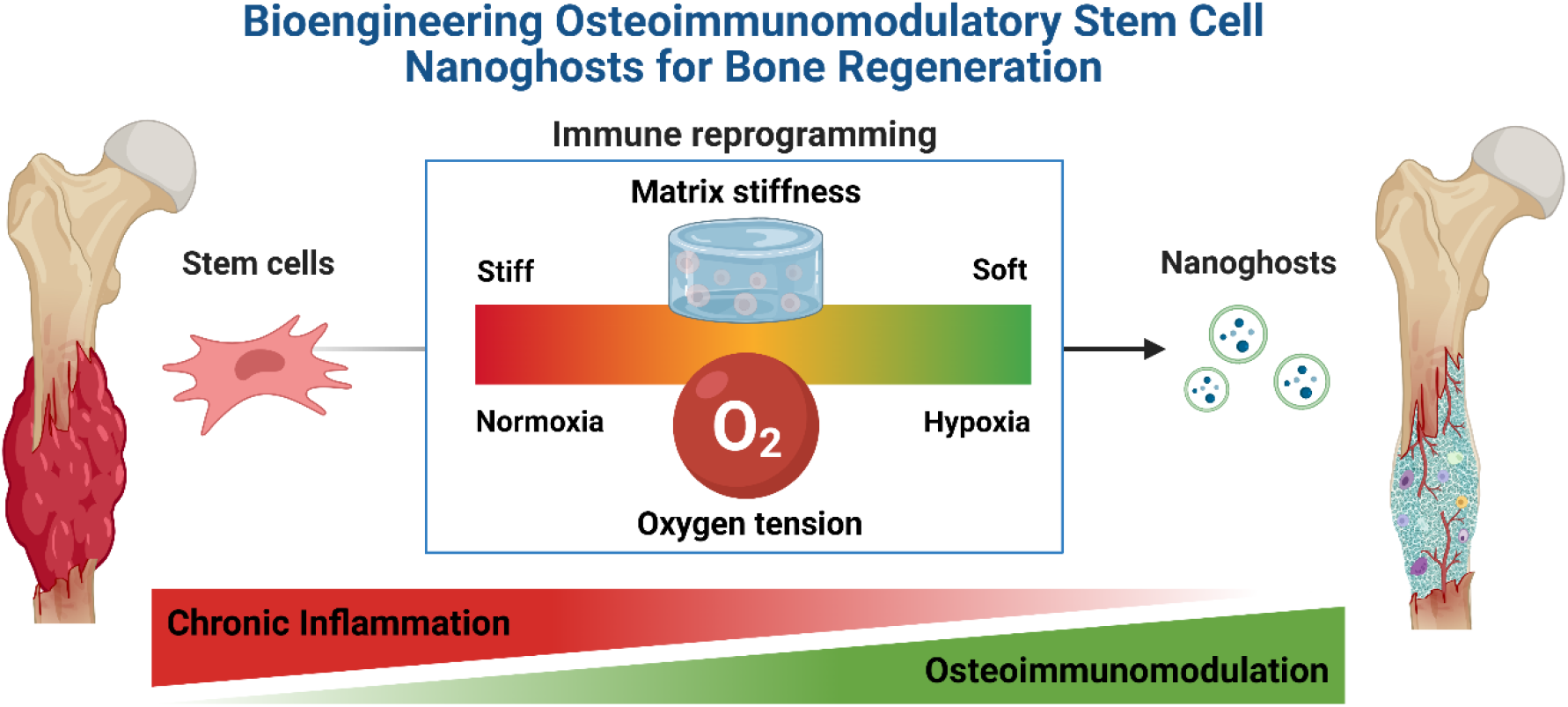

## 1. Introduction

Bone fractures arising from trauma, osteoporosis, or tumor resection constitute a growing global health challenge, with critical-sized defects often necessitating surgical intervention ^1,2^. In 2019, an estimated 178 million new fractures were reported worldwide, a burden that continues to increase with population ageing ^3^. The clinical and socioeconomic impact is substantial: osteoporotic fractures alone account for up to €57 billion in annual healthcare expenditures in Europe and are a leading cause of long-term disability in older adults ^4^. Current management of bone fractures relies on surgical reconstruction using autologous bone grafts and biomedical implants, including titanium screws, plates, and prostheses ^5^. While essential for mechanical stabilization, clinical outcomes remain suboptimal, with implant failure rates of 5-15% in complex repairs and substantially higher rates in patients with impaired bone healing ^6,7^. These failures are frequently driven by dysregulated inflammatory responses at the injury-implant interface, which hinder osseointegration, promote fibrotic encapsulation, and compromise long-term stability, often necessitating revision surgery ^8,9^. Collectively, these challenges underscore an urgent need for regenerative strategies that not only promote bone formation but also actively modulate the local immune microenvironment to improve implant integration and durable repair.

Recent advances in osteoimmunology have revealed that bone regeneration is not solely a process of osteogenesis but is critically governed by dynamic interactions between skeletal and immune systems ^10,11^. A tightly regulated inflammatory response is essential during the early phases of healing, facilitating debris clearance, progenitor cell recruitment, angiogenesis, and matrix remodeling ^12^. However, persistent or dysregulated inflammation can be detrimental. Conventional orthopedic implants and synthetic biomaterials, such as titanium screws, fixation plates, and spinal cages, often elicit foreign body responses that can sustain chronic inflammation, impairing osseointegration and lead to fibrosis, implant loosening, or failure ^8,9^. In this context, immunoregulatory therapies have emerged as a promising paradigm for bone tissue engineering, aiming to actively shape the local immune microenvironment to support regenerative outcomes.

Human mesenchymal stromal/stem cells (hMSCs) play a central role in bone repair due to their dual capacity to differentiate into osteogenic lineages and to modulate immune responses ^13,14^. Nevertheless, the clinical translation of hMSCs has been hampered by low engraftment efficiency, high manufacturing costs, and complex regulatory pathways ^15,16^. Mounting evidence indicates that the regenerative efficacy of hMSCs is predominantly mediated by paracrine mechanisms rather than direct cell replacement ^17^. In particular, extracellular vesicles (EVs), nanoscale particles enriched in bioactive proteins, lipids, and nucleic acids, have been shown to recapitulate many of the biological effects of their parental cells ^18–20^. As acellular agents, EVs offer advantages in safety, storage, and immunogenicity, positioning these vesicles as an attractive alternative to cell-based therapies ^21,22^. However, the translation of EV-based approaches is challenged by low yields, heterogeneity, and difficulties in large-scale manufacturing and standardization ^21,23^. To overcome these limitations, nanoghosts (NGs) or cell-derived nanovesicles have emerged as a compelling EV-mimetic platform. NGs are plasma-membrane-derived nanoparticles generated through controlled physical disruption and reassembly of cell membranes, resulting in nano-sized vesicles that retain key surface proteins and molecular signatures of their parental cells ^24,25^. Compared to naturally secreted EVs, NGs offer significantly higher nanoparticle yield, shorter processing times, greater scalability, and enhanced versatility for cargo loading ^26,27^. Moreover, NG production overcomes the cargo-selection constraints inherent to EV biogenesis. Thus, by decoupling therapeutic function from living cells, these NGs retain the benefits of cell therapies; while offering greater safety, reproducibility, off-the-shelf-use, and scalability needed for clinical translation.

Given the plasticity of hMSCs in response to biophysical and biochemical cues, such as substrate stiffness, oxygen tension, mechanical loading, and 3D architecture ^28–30^, there is growing interest in exploiting physiological conditions to enhance cellular functionality. Conventional 2D tissue culture substrates are supraphysiologically stiff (Young’s modulus ∼3 GPa), far exceeding that of native tissues (∼0.1 kPa - 1 MPa), which can compromise MSC phenotype ^31,32^. For instance, MSCs cultured on softer matrices (∼3 kPa) produce higher levels of anti-inflammatory cytokines (i.e. IL-10, CCL17), promoting anti-inflammatory macrophage polarization and attenuating inflammation *in vivo* when compared to cells cultured in stiff matrices (∼30 kPa)^33^. Similarly, oxygen tension profoundly influences MSC function: bone marrow-derived MSCs naturally reside in hypoxic niches (1 - 7% O_2_), and culture under low oxygen enhances proliferation, differentiation potential, and immunomodulatory activity relative to atmospheric conditions ^34,35^. Harnessing such biomimetic cues could provide a strategy to engineer osteoimmunomodulatory NGs capable of orchestrating immune– bone crosstalk and enhancing bone regeneration.

In this study, we showcase the bioengineering of hMSC-derived osteoimmunomodulatory NGs as a scalable, acellular platform for bone repair. Initially, two NG methods were compared to evaluate nanoparticle production yield - spin cup (SC) and freeze-thaw (FT) (Fig. 1). Next, the reprogramming of hMSCs immunophenotype within biomimetic microenvironments were evaluated, specifically tuning oxygen tension and matrix stiffness. Finally, the osteoimmunomodulatory capacity of hMSC-NGs in an *in vitro* inflammatory model was evaluated, establishing a foundation for their application in regenerative bone therapy.

**Figure 1.**
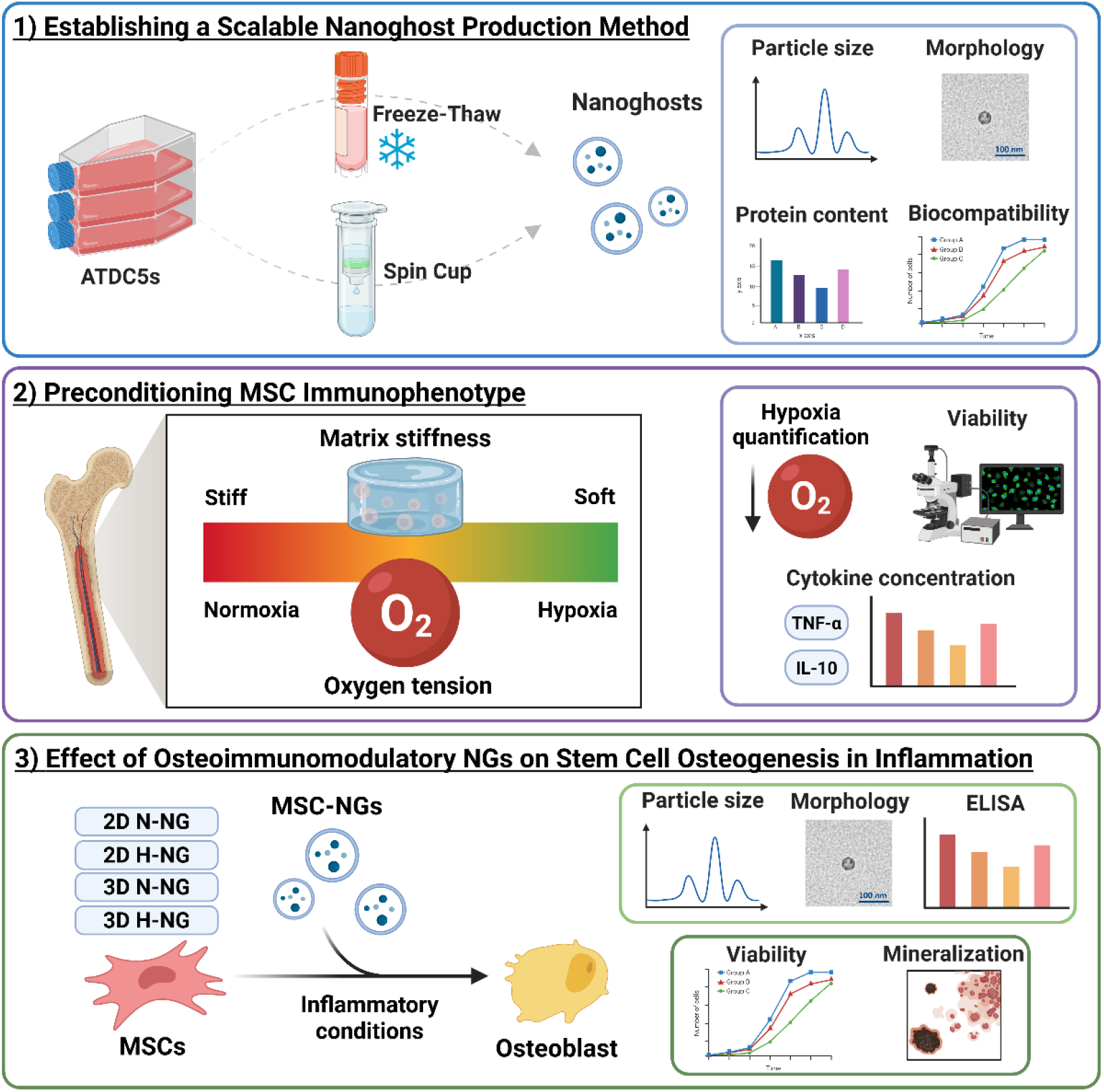
Experimental workflow for the generation and evaluation of osteoimmunomodulatory NGs. 1) Establishing a scalable NG production method. ATDC5 cells are processed using FT and SC methods to generate NGs. NGs are characterized by particle size distribution, morphology, protein content, and biocompatibility. 2) Preconditioning MSC immunophenotype. hMSCs are cultured under defined biophysical and biochemical conditions by varying matrix stiffness (stiff to soft) and oxygen tension (hypoxia to normoxia) to modulate their immunophenotype. Hypoxia levels, cell viability, and cytokine secretion (e.g., TNF-α and IL-10) are quantified to assess immunomodulatory responses. 3) Effect of osteoimmunomodulatory NGs on hMSC osteogenesis under inflammatory conditions. NGs derived from hMSCs cultured in 2D or 3D environments under normoxic (N) or hypoxic (H) conditions (2D N-NG, 2D H-NG, 3D N-NG, 3D H-NG) are applied to hMSCs exposed to inflammatory cues. hMSC-NGs are characterized for size, morphology and cytokine content, and their effects on recipient hMSCs viability and osteogenic differentiation under inflammatory conditions are assessed. Figure created with biorender.com.

## 2. Methods

### 2.1. Cell culture

The ATDC5 chondrogenic cell line was maintained in a basal medium consisting of DMEM/F-12 (Gibco, 11320033) supplemented with 5% heat-inactivated fetal bovine serum (FBS, S14068S1810, Biowest), and 100 U/mL penicillin/100 µg/mL streptomycin (Gibco, 15140). Cells were maintained at 37°C under humidified conditions and 5% carbon dioxide (CO_2_), and the medium was replaced twice a week. Cells were passaged at 80% confluence.

hMSCs were isolated from bone marrow aspirates of healthy patients (N=2, 12 and 20 years of age, female) after informed consent, in accordance with a biobank protocol approved by the local Medical Ethics Committee (TCBio-08-001, University Medical Center Utrecht) ^36^. Adherent cells were maintained at 37°C under humidified conditions and 5% CO_2_ in hMSCs basal medium consisting of α-MEM (22 561, Invitrogen) supplemented with 10% heat-inactivated FBS, 0.2 mM L-ascorbic acid 2-phosphate (A8960, Sigma-Aldrich), 100 U/mL penicillin with 100 mg/mL streptomycin, and 1 ng/mL basic fibroblast growth factor (233-FB; R&D Systems). The media were refreshed every 2 days until the cells reached 80% confluence. All experiments described in this study using hMSCs were conducted at passage 4.

### 2.2. NGs production

#### 2.2.1. Freeze-thaw (FT) method

NGs were produced via the FT method as described previously ^37^. ATDC5s (1 × 10^7^ cells) were resuspended in 1 mL deionized H_2_O. Samples were incubated for 30 min at room temperature, followed by three FT cycles between -80°C and 37°C. After processing, samples were centrifuged at 2,000 g for 20 min to remove debris. The supernatant was collected for purification by ultracentrifugation.

#### 2.2.2. Spin cup (SC) method

NGs were produced via the SC method as described previously ^26,38^. ATDC5s (1 × 10^7^ cells) were resuspended in 300 µL PBS. The suspension was loaded into a spin cup containing a 10 µm filter (69705, Sigma-Aldrich) and centrifuged at 14,000 g for 10 min at 4°C. The flow-through was collected and centrifuged a second time under the same conditions. The flow-through was then transferred to a spin cup fitted with an 8 µm filter (TETP04700, Sigma-Aldrich) and centrifuged twice at 14,000 g for 10 min at 4°C. The final flow-through was collected for purification by ultracentrifugation.

### 2.3. NG isolation and characterization

NGs were isolated using an Optima XE-90 ultracentrifuge with a SW 32 Ti rotor (Beckman Coulter, USA). Samples were centrifuged at 10,000 g for 30 mins to obtain the large NG population, whilst the supernatant was spun at 120,000 g for 70 mins. The NG pellet was washed with fresh PBS and spun for 120,000 g for 70 mins. The remaining NG pellet was combined with the large NG population and stored at -80°C until required.

Protein concentrations of NGs were quantified using the MicroBCA Protein Assay kit (Thermo Scientific) according to the manufacturer’s instructions. Nanoparticle tracking analysis (NTA) was conducted to evaluate particle size distribution and concentration using the ZetaView® Evolution (Particle Metrix GmbH).

Transmission electron microcopy (TEM) of isolated NGs was achieved using the TFS Tecnai 20 transmission electron microscope. Samples were physisorbed to 200 µm mesh, carbon-coated copper formvar grids (Agar Scientific) and negatively stained with 1% uranyl acetate.

To compare NG and EV yield, hMSCs (2 × 10^6^ cells) were cultured for 48 h in basal medium containing EV-free FBS. The conditioned medium was collected and EVs were isolated with differential ultracentrifugation as previously described ^20^. The hMSCs were processed for NG production as described above.

### 2.4. NG biocompatibility

ATDC5 cells were seeded at 1 × 10^3^ cells/well in 96-well plates. After 24 h, medium was replaced with fresh basal medium supplemented with SC- and FT-NGs (1 × 10^7^ particles per well) and cultured under standard conditions (37°C, 5% CO_2_) for up to 7 days. One plate per time point (days 1, 3, 7) was analyzed. Viability was assessed using the Alamar Blue Assay (R7017, Sigma-Aldrich) by replacing the medium with fresh medium containing 10% v/v Alamar Blue and incubating for 4 h at 37°C. Fluorescence was recorded at an excitation of 560 nm and an emission of 590 nm. Following which, cells were lysed with 0.1% Triton X-100 in PBS and then stored at -80°C. DNA content was quantified using the Quant-iT PicoGreen dsDNA Assay Kit (BP7589, Invitrogen), measuring fluorescence at an excitation of 485 nm and an emission of 520 nm.

### 2.5 hMSC preconditioning and characterization

#### 2.5.1. Oxygen tension preconditioning

hMSCs were cultured in T175 flasks for 48 h under normoxia (21% O_2_) and hypoxia (5% O_2_). Following incubation, conditioned medium was collected for downstream cytokine analysis, whilst the hMSCs (2 × 10^6^ cells) were harvested for NG production. The effect of oxygen tension conditioning on hypoxia in cells was evaluated using the Image-iT™ Green Hypoxia Reagent (Thermo Fisher Scientific, I14833) following manufacturer’s guidelines. Cultures were incubated with a 5 µM working solution with complete medium (4 h in the dark) under their respective oxygen conditions. After being washed in complete medium, fluorescence intensity (Ex/Em 480/520 nm) was measured using the CLARIOstar Plus microplate reader (BMG Labtech).

### 2.6. 3D substrate preconditioning

#### 2.6.1. GelMA Synthesis

GelMA was synthetized as previously described ^39^. Briefly, type A porcine skin gelatin (Sigma-Aldrich) was dissolved at 10% (wt/vol) in PBS. Methacrylic anhydride (Sigma-Aldrich) was then added at 0.6 g/mL to the gelatin solution, which was continuously stirred at 50 °C for 1 h. The resulting mixture was dialyzed against distilled water using 12-14 kDa cutoff tubing (Thermo Scientific) for 2-3 days at 40°C to remove residual salts and methacrylic acid. Post-dialysis, the GelMA solution was diluted to 2% (w/v) and the pH adjusted to 7.4 with 1 mM sodium hydroxide (Sigma-Aldrich). The solution was subsequently sterilized via filtration and lyophilized over 2 days. The lyophilized GelMA was stored at –80°C until further use.

#### 2.6.2. Fabrication of hMSC-GelMA constructs

hMSCs were encapsulated in GelMA hydrogels (1.5 × 10^6^ cells/mL) at a final concentration of 5% and 15% (w/v) using visible-light photoinitiators (1 mM Tris (2,2′-bipyridyl) dichlororuthenium (II) hexahydrate (Ru) (Sigma-Aldrich) and 10 mM Sodium persulfate (SPS) (Sigma-Aldrich)). Hydrogels were cast as 200-µL constructs in 24-well suspension plates. Photocrosslinking was performed (10 min, RT) under visible light (30 mW/cm^2^). After crosslinking, constructs were washed with basal MSC medium and cultured 48 h under normoxic or hypoxic conditions. Conditioned medium was collected for downstream cytokine analysis. After conditioning, hMSCs were retrieved using Collagenase Type II (Sigma-Aldrich, Cat. No. C6885) at 250 U/mL in PBS (37°C) for 30 mins. Cell suspensions were passed through a 70-µm cell strainer, pelleted (300 g, 5 min), and hMSCs (2 × 10^6^ cells) were harvested for NG production.

#### 2.6.3. Cellular morphology within GelMA constructs

hMSC morphology within the 5 wt-% and 15 wt-% GelMA hydrogels was assessed using a Live/Dead Calcein-AM/Ethidium homodimer-1 (EthD-1) staining as previously described ^40^. Constructs were imaged using confocal microscopy (Leica Microsystems, Germany). Z-stacks were converted into maximum-intensity projections prior to analysis. Quantitative analysis of average cell surface area (calcein positive cells) was performed using ImageJ (NIH). Three random locations were selected per sample.

### 2.7. Inflammatory cytokine assessment

Concentrations of pro-inflammatory cytokine TNF-α and anti-inflammatory cytokine IL-10 within the collected hMSCs conditioned medium and hMSC-derived NGs were quantified using commercial ELISA kits (R&D Systems, DTA00D and D1000B), according to the manufacturer’s instructions. NGs were lysed with Triton X-100 (0.1%, Sigma-Aldrich) with several freeze/thaw cycles prior to use. Absorbance was read at 450 nm using the CLARIOstar Plus microplate reader (BMG Labtech). Concentrations were calculated using appropriate standard curves. Medium alone and 0.1% Triton X-100 was used as the background for the condition medium and NGs, respectively.

### 2.8. Cell uptake assay for hMSC-NGs

hMSCs uptake was evaluated using CellMask-labelled NGs as described previously ^41^. In brief, NGs were stained with CellMask™ Deep Red Plasma Membrane Stain (5 μg/mL in PBS; Thermo Scientific) for 10 minutes, followed by two washing steps in PBS using ultracentrifugation at 120,000 g for 70 minutes. hMSCs were seeded at a density of 4 × 10^3^ cells/cm^2^ for 24 h. The culture medium was then replaced with fresh basal medium containing labelled NGs, and cells were incubated for 8 h. Following incubation, cells were fixed with neutral buffered formalin and permeabilized with triton-X. The actin cytoskeleton was labelled with Alexa Fluor 488 phalloidin (1:20) (Cell Signalling Technology) and then mounted with Prolong ™ Gold Antifade Mountant with DAPI (Thermo Scientific) to label the nuclei. Imaging was performed using confocal laser scanning microscopy (Leica Microsystems, Germany).

### 2.9. Metabolic activity of NG treated hMSCs under inflammatory conditions

hMSCs were seeded in 96-well plates at a density of 3 × 10^3^ cells/cm^2^ in basal medium. After 24 h, medium was replaced with basal medium supplemented with NGs (1 µg/ml) and either TNF-α (10 ng/mL) or IL-1β (10 ng/mL). Metabolic activity was assessed on days 1, 3 and 7 using the Alamar Blue reagent. Medium was replaced at every timepoint. Cells cultured in cytokine-supplemented basal medium without NGs were used as the control.

### 2.10. Osteogenic capacity of NG treated hMSCs under inflammatory conditions

hMSCs were seeded in 96-well plates at a density of 21 × 10^3^ cells/cm^2^ in basal MSC medium. After 24 h, medium was replaced with osteogenic medium (consisting of basal medium with 10 mM β-glycerophosphate and 10 nM dexamethasone) supplemented with NGs (1 µg/ml) and either TNF-α (10 ng/mL) or IL-1β (10 ng/mL). After 7 days, cells were cultured in cytokine-free osteogenic medium supplemented with NGs (1 µg/ml) until day 21. Medium was changed every 2 days. Cells cultured in osteogenic medium or cytokine-supplemented osteogenic medium were used as the controls.

### 2.11. Calcium Deposition and Mineralization

To evaluate hMSC mineralization capacity, Alizarin red S staining for calcium production was conducted as previously described ^41^. Samples were washed with PBS and fixed in 10% formalin for 30 min. Next, samples were washed with distilled water and incubated with 40 mM Alizarin Red S solution (pH 4.1, Sigma Aldrich) for 10 min. The unbound staining solution was removed by distilled water washes. Staining was visualized using light microscopy (Olympus Microscope BX43). For alizarin red quantification, samples were de-stained with 10% cetylpyridinium chloride (Sigma-Aldrich) for 1 h and the absorbance was read at 550 nm using the CLARIOstar Plus microplate reader (BMG Labtech).

### 2.12. Statistical analysis

For all data presented, experiments were repeated at least 3 times. All statistical analysis was undertaken using ANOVA multiple comparisons test with Tukey modification (GraphPad Prism 8). P values equal to or lower than 0.05 were considered significant. *P ≤ 0.05, **P ≤ 0.01, ***P ≤ 0.001, and ****P ≤ 0.0001.

## 3. Results

### 3.1. Establishing a Reproducible and Scalable NG Production Method

The ATDC5 cell line was used to assess the two NG methods (SC and FT) as they provide a reproducible model not restricted by the donor-to-donor variability and limited expansion capacity of primary cells. SC- and FT-derived NGs displayed a similar size distribution and vesicle-like morphology, as demonstrated by NTA and TEM (Fig. 2A). Both NG groups exhibited similar particle sizes within the nanometer range (P > 0.05) (Fig. 2C). Notably, FT-NGs were produced at a significantly higher concentration (426-fold) (∼16,000 particles/cell) than SC-NGs (∼377 particles/cell) (Fig. 2C) (P ≤ 0.01), which was consistent with their increased protein content (Fig. 2D) (P ≤ 0.01).

**Figure 2.**
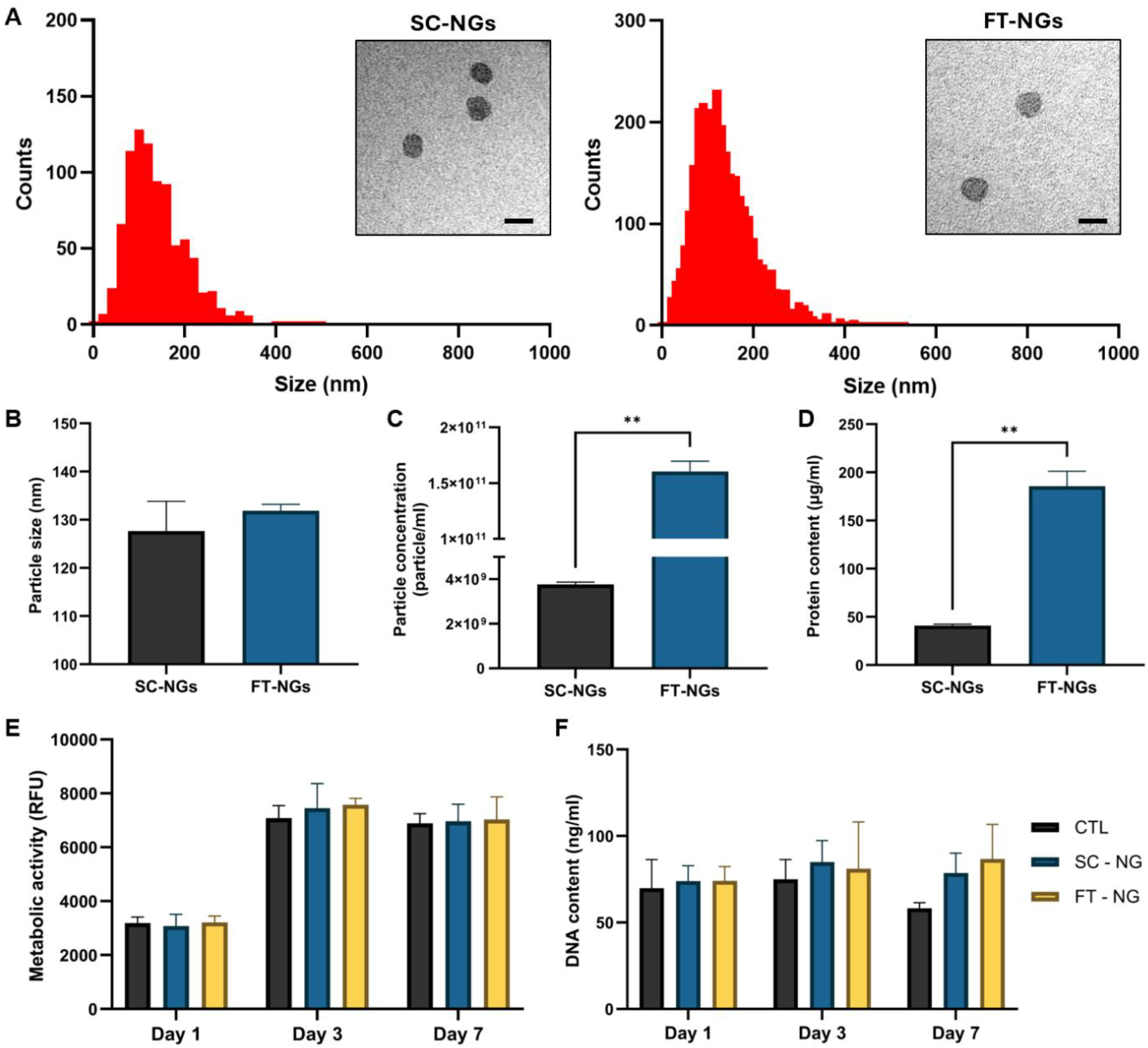
Validation of NG manufacturing methods and biocompatibility. ATDC5 NGs created using the SC and FT methods were assessed in terms of their A) size distribution and morphology (Scale bar = 100 nm.), B) average particle size, C) particle concentration, and D) protein content. NG biocompatibility on ATDC5 cells were evaluated by assessing E) metabolic activity, and F) DNA content over 7 days. Data expressed as mean ± SD. **P ≤ 0.01.

The biocompatibility of the generated NGs was evaluated by assessing ATDC5 metabolic activity and DNA content. Both SC- and FT-NGs exhibited comparable biocompatibility to the untreated control over 7 days of culture, with no significant differences observed in metabolic activity or DNA content (Fig. 2E, F) (P > 0.05).

### 3.2. Biomimetic Microenvironments Modulate hMSCs Immunophenotype

The influence of biomimetic microenvironmental cues, specifically matrix stiffness and oxygen tension, on modulating the immunomodulatory phenotype of hMSCs was examined. The influence of matrix stiffness using GelMA substrates was evaluated. hMSCs maintained high viability after 48 h of culture within both 5 wt% and 15 wt% GelMA hydrogels (Fig. 3A). Cells cultured within 5 wt% GelMA exhibited a slight increase in cell surface area relative to those in 15 wt% GelMA (Fig. 3B) (P > 0.05). Analysis of cytokine secretion revealed that hMSCs cultured within the stiffer 15 wt% GelMA secreted significantly higher levels of TNF-α compared with those in the 5 wt% GelMA condition (1.95-fold) (P ≤ 0.001), whereas IL-10 secretion was significantly elevated in hMSCs cultured within the softer 5 wt% GelMA matrix (1.45-fold) (Fig. 3C) (P ≤ 0.001).

**Figure 3.**
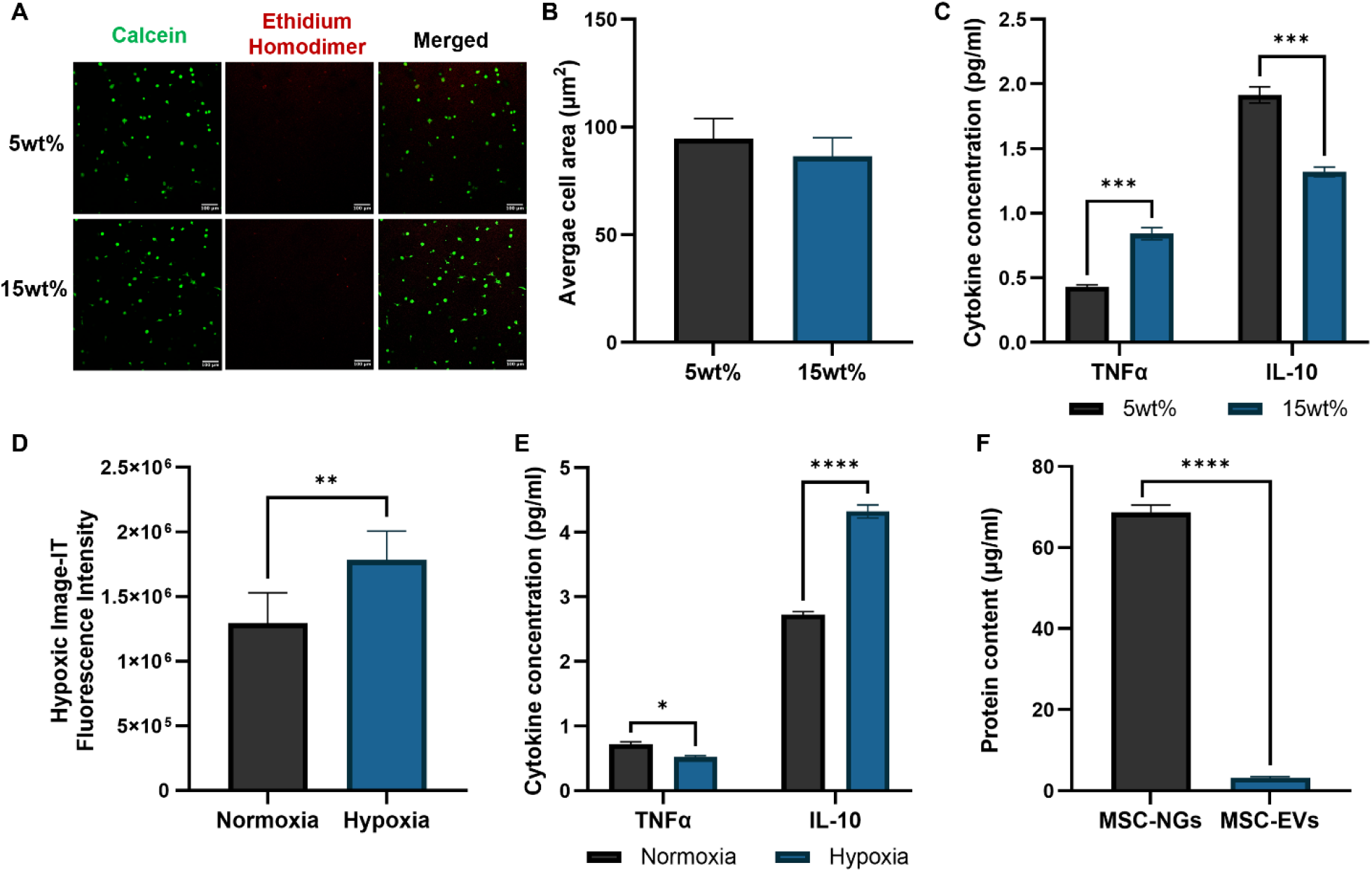
hMSC preconditioning via altering substrate stiffness and oxygen tension. A) Live/dead staining of 5 or 15 wt% GelMA encapsulated hMSCs. B) Average cell surface area of hMSCs within 5 or 15wt% GelMA. C) Cytokine analysis of collected conditioned medium from hMSCs within 5 or 15wt% GelMA. D) Hypoxic image-IT fluorescence analysis of normoxia or hypoxia-treated hMSCs. E) Cytokine analysis of collected conditioned medium from normoxic or hypoxic hMSCs. F) Protein content of hMSC-derived NGs and EVs. Data expressed as mean ± SD. *P ≤ 0.05, **P ≤ 0.01, ***P ≤ 0.001 and ****P ≤ 0.0001.

hMSCs were preconditioned under either normoxic or hypoxic conditions for 48 h. hMSCs exposed to hypoxia exhibited significantly higher Hypoxic Image-IT fluorescence intensity when compared to cells in normoxia (Fig. 3D) (P ≤ 0.01). To evaluate the impact of oxygen tension on immunomodulatory function, inflammatory cytokines in the conditioned medium were quantified. Hypoxic preconditioning significantly reduced TNF-α secretion compared with normoxic culture (1.4-fold) (P ≤ 0.05), while concomitantly increasing IL-10 levels (1.59-fold) (Fig. 3E) (P ≤ 0.0001). The comparative yields of NGs and EVs produced from hMSCs were evaluated. Our findings showed that a significantly higher quantity of NGs (>21.6-fold) were produced compared to EVs from the same cells (Fig. 3F).

### 3.3. Osteoimmunomodulatory NGs Protect hMSCs Viability within an Inflammatory Environment

Following hMSC preconditioning through modulation of oxygen tension (normoxia and hypoxia) and substrate stiffness (2D and 3D (5wt% GelMA)), NGs were generated using the FT method. All NG groups exhibited nanoscale particle size distributions and vesicle-like morphology, as confirmed by NTA and TEM (Fig. 4A), indicating that preconditioning did not alter fundamental NG structural characteristics.

**Figure 4.**
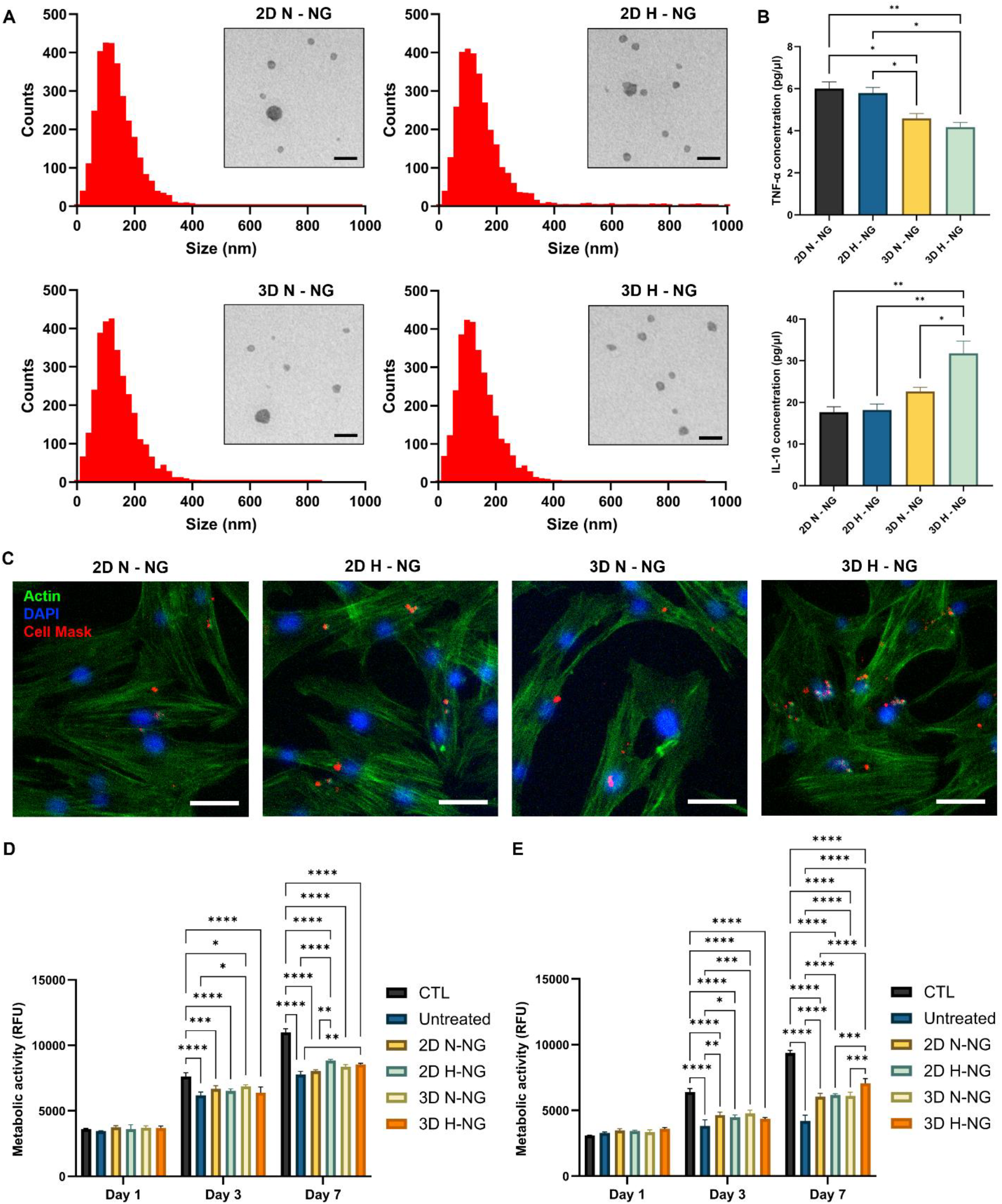
The effect of hMSC-NGs on hMSCs viability under inflammatory conditions. A) NTA and TEM analysis of hMSC NGs. Scale bar = 100 nm. B) TNF-α and IL-10 concentration of hMSC NGs. D) NG uptake by hMSCs following 8 hours incubation. Scale bar = 50 µm. Biocompatibility of hMSCs treated with/without NGs under D) IL-1β or E) TNFα exposure. Data expressed as mean ± SD. *P ≤ 0.05 and **P ≤ 0.01.

To determine whether microenvironmental preconditioning influenced hMSC-NG immunological cargo, TNF-α and IL-10 content was quantified. Our findings showed that hMSC-NGs generated under hypoxia combined with 3D conditions (3D H-NGs) contained lower TNF-α levels when compared to NGs obtained from 3D normoxic, 2D hypoxic (P ≤ 0.05), and 2D normoxic conditions (P ≤ 0.01) (Fig. 4B). Moreover, NGs from hMSCs cultured under combined hypoxic and 3D conditions (3D H-NGs) exhibited significantly elevated IL-10 levels compared with NGs derived from 3D normoxic (P ≤ 0.05), 2D hypoxic (P ≤ 0.01), and 2D normoxic cultures (P ≤ 0.01) (Fig. 4B). These findings suggest that integration of hypoxic and 3D microenvironmental cues synergistically enhances the immunomodulatory payload of hMSC-derived NGs.

Next the biological function of these NGs was assessed by evaluating their uptake by recipient hMSCs (Fig 4C). Our findings showed that NGs from all groups were internalized by hMSCs after 8 hours of incubation. The capacity of hMSC-derived NGs to protect hMSC viability under inflammatory stress was explored. Following treatment with IL-1β or TNF-α, hMSCs viability was significantly reduced on day 3 and 7 compared to the control (P ≤ 0.0001) (Fig 4D, E). Moreover, NGs treatment was shown to increase the metabolic activity of IL-1β or TNF-α exposed hMSCs on day 3 and 7. Specifically, under IL-1β exposure, 3D N-NG exhibited a significant increase in metabolic activity compared to the untreated control on day 3 (P ≤ 0.05), whilst 2D H-NG and 3D H-NG displayed increased metabolic activity compared to the untreated control on day 7 (P ≤ 0.01) (Fig 4D). Under TNF-α exposure, 2D N-NG (P ≤ 0.01), 2D H-NG (P ≤ 0.05), and 3D N-NG (P ≤ 0.001) displayed a significant increase in metabolic activity compared to the untreated control on day 3, whilst all NG-treated groups demonstrated significantly improved metabolic activity when compared to the NG-free control (P ≤0.0001), with 3D H-NGs conferring the greatest protective effect (Fig. 4E). Collectively, these results indicate that hMSC preconditioning within hypoxic and 3D microenvironments enhances the functional efficacy of derived NGs, promoting cell viability under inflammatory conditions.

### 3.4. Osteoimmunomodulatory NGs Improve hMSCs Mineralization Capacity within an Inflammatory Environment

To assess whether hMSC-derived NGs could rescue hMSCs osteogenic capacity under inflammatory conditions, calcium deposition was evaluated following 21 days of osteogenic induction in the presence of IL-1β or TNF-α (Fig. 5). Alizarin Red S staining revealed a pronounced reduction in mineralized matrix formation in cytokine-treated hMSCs compared with untreated controls, confirming inflammation-mediated suppression of osteogenesis (Fig. 5A).

**Figure 5.**
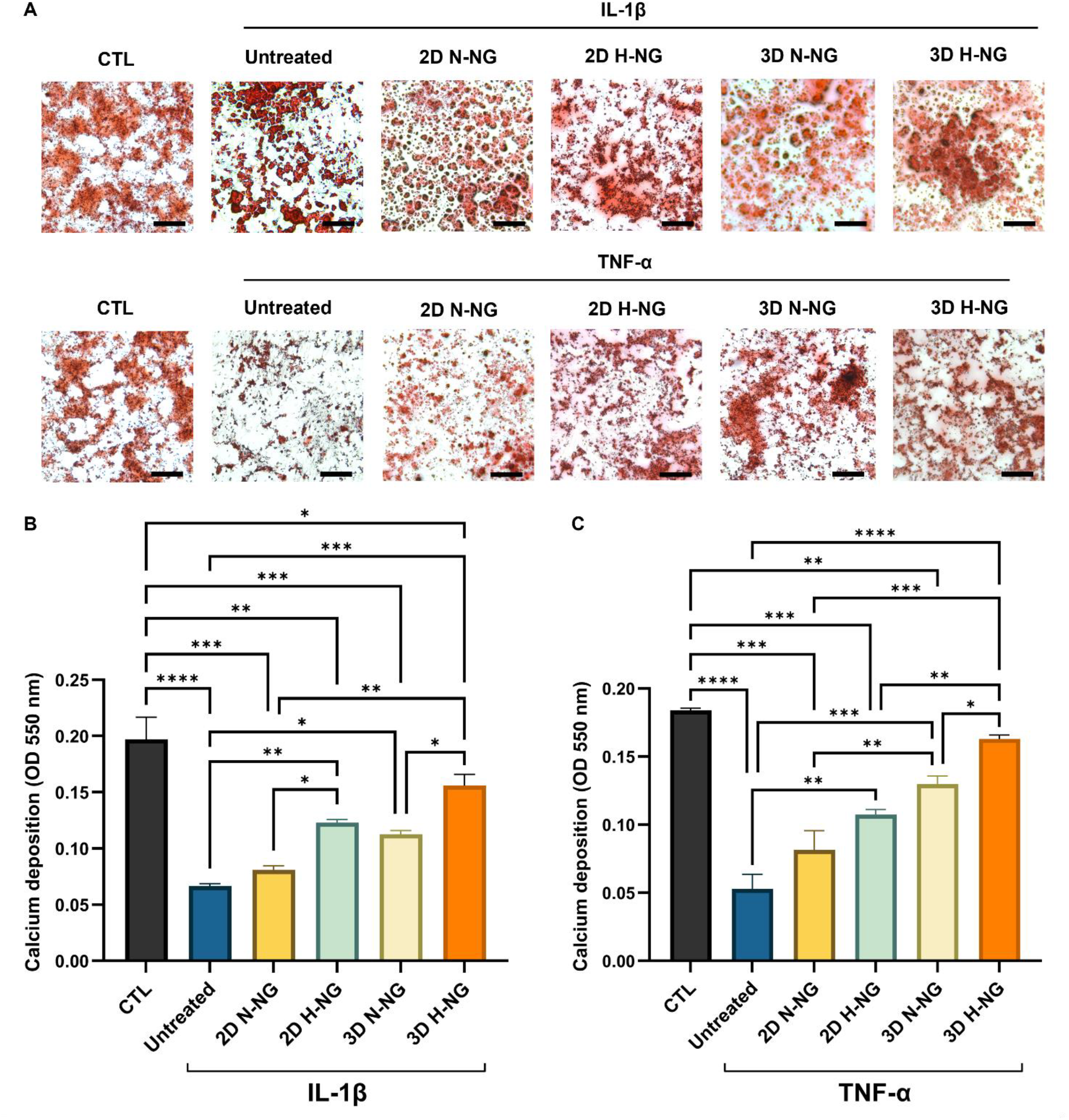
The effect of hMSC-NGs on the osteogenic differentiation of hMSCs under inflammatory conditions. A) Alizarin red S staining for calcium deposition of hMSC-NG treated hMSCs under inflammatory conditions following 21 days of osteogenic culture, including an untreated control (without NGs) and a control (CTL) without inflammatory stimuli present. Scale bars = 200 µm. Quantitative analysis of calcium deposition of the NG treated hMSC groups as under A is shown for B) IL-1β or C) TNF-α induced inflammation. Data expressed as mean ± SD. *P ≤ 0.05, **P ≤ 0.01,***P ≤ 0.001 and ****P ≤ 0.0001.

Quantitative analysis demonstrated that IL-1β exposure significantly reduced calcium deposition relative to the control group (Fig. 5B) (P ≤ 0.0001). Treatment with hMSC-derived NGs significantly increased mineralization across all NG conditions (except 2D N-NGs) compared with the cytokine-only control (P ≤ 0.05 - 0.001). Notably, NGs derived from hypoxia-conditioned hMSCs promoted greater calcium deposition than those derived under normoxic conditions, with 3D hypoxic NGs (3D H-NGs) exhibiting the highest mineralization levels among all NG-treated groups. A similar trend was observed under TNF-α challenge (Fig. 5C). TNF-α treatment markedly impaired calcium deposition compared to the cytokine-free control (P ≤ 0.0001), whereas NG administration significantly restored mineralization in a preconditioning-dependent manner. 3D H-NGs induced the greatest recovery of calcium deposition, approaching levels observed in untreated controls (P > 0.05). Collectively, these results demonstrate that hMSC-derived NGs partially protected inflammation-induced inhibition of osteogenic differentiation, with NGs generated under combined hypoxic and 3D microenvironmental conditions exhibiting superior osteogenic rescue.

## 4. Discussion

Bone regeneration is a tightly orchestrated process governed by reciprocal interactions between skeletal and immune systems ^10,42^. While early inflammatory cues are essential for initiating repair, sustained or dysregulated inflammation severely compromise osteogenesis and remain a major cause of non-union and implant failure ^22,43^. In this study, we establish bioengineered hMSC-derived NGs as a scalable, acellular nanotherapeutic platform capable of actively modulating osteoimmune crosstalk. By combining biomimetic hMSC preconditioning with a high-yield NG manufacturing, out findings demonstrate that NGs can be endowed with tailored immunoregulatory cargo and functional efficacy, ultimately restoring osteogenic differentiation under inflammatory stress.

A critical barrier to the translation of EV-based therapies has been limited production yield and batch-to-batch heterogeneity ^21,22,44^. Recent research has demonstrated the increased particle yields generated by NG methods when compared to EV isolation ^26,27^, however, there has been a lack of studies in the field directly comparing the production efficiency of different NG methods. Our comparison of established NG fabrication strategies revealed that the FT method produces significantly higher nanoparticle yields compared to the SC approach without compromising biocompatibility. Importantly, both methods preserved nanoscale size and vesicle-like morphology, consistent with findings in the literature ^26,37^. Moreover, the enhanced protein content observed in the FT-NGs further suggests improved retention of bioactive components, which may underlie their functional potency. Critically, we demonstrated superior hMSCs NG yield when compared to EVs secreted from the same cells, thus supporting FT-derived NGs as a scalable and reproducible alternative to naturally secreted EVs, addressing a central limitation in current acellular regenerative strategies.

Beyond scalable production, a key aspect of this research was to demonstrate that NG bioactivity can be rationally programmed through microenvironmental MSC conditioning. There has been increasing research employing physiological stimuli (i.e. hypoxia, 3D architecture, matrix stiffness) to reprogram cell function ^45^. For instance, researchers have harnessed hypoxic preconditioning to improve the osteogenic capacity of MSCs and their secretome ^28,46^. Moreover, matrix stiffness and architecture have been shown to modulate the differentiation capacity of cells ^29,47^. Our findings showed that hypoxic preconditioning suppresses pro-inflammatory TNF-α secretion while enhancing IL-10 release, consistent with hypoxia-driven immunoregulatory reprogramming reported in MSCs ^48^. Similarly, culture within softer GelMA matrices promoted an anti-inflammatory secretory profile relative to stiffer substrates, aligning with reports that softer matrices shift MSCs toward an anti-inflammatory phenotype ^33^. Notably, these immunomodulatory shifts occurred without compromising viability, indicating phenotypic reprogramming rather than stress-induced effects.

Strikingly, these microenvironmental alterations to MSCs phenotype were inherited by the derived NGs. hMSC-NGs generated under combined hypoxic and 3D conditions exhibited markedly elevated IL-10 content, indicating that NGs capture not only the membrane architecture ^49^, but also the functional immunological cargo of their parental cells. These findings support the concept that NGs act as “phenotypic snapshots” of conditioned cells, offering a powerful means to decouple therapeutic function from living cell delivery. Moreover, the capacity of NGs to functionally deliver cellular contents overcome issues associated with EVs, where vesicle-associated cargo is inherently dictated by cell type and EV biogenesis route, limiting control over what therapeutic cargo is enriched within vesicles without complex cell engineering ^50,51^. Thus, NGs may provide a better platform to deliver endogenous cell-derived bioactive factors when compared to EV-based systems. The synergistic enhancement observed under combined hypoxic and 3D conditions further underscores the importance of integrating biochemical and biophysical cues when engineering next-generation nanotherapeutics.

Both IL-1β and TNF-α are well-established inhibitors of MSC viability and osteogenic differentiation, recapitulating pathological conditions observed in non-healing fractures ^52,53^. Our findings showed that NG treatment preserved hMSCs viability under inflammatory conditions, with hypoxia-derived NGs, and particularly 3D H-NGs, exerting the strongest protective effects. These results suggest that NGs buffer inflammatory stress, potentially by delivering anti-inflammatory mediators such as IL-10 or by modulating receptor-level signaling at the cell membrane. Whilst the precise molecular mechanisms warrant further investigation, the observed functional rescue highlights NGs as active immunoregulatory agents rather than passive delivery vehicles.

Crucially, this immunomodulatory protection translated into restored osteogenic function. Inflammatory cytokine exposure profoundly impaired long-term hMSCs mineralization, consistent with inflammation-mediated suppression of osteogenic transcriptional programs ^54,55^. Treatment with hMSC-derived NGs partially rescued calcium deposition in both IL-1β- and TNF-α-challenged cultures, with NGs derived from hypoxic 3D hMSCs consistently yielding the greatest improvement. These results reinforce the concept that effective bone regeneration requires not only osteoinductive cues but also active immune modulation. By mitigating inflammatory inhibition, NGs enable MSCs to re-engage osteogenic pathways, thereby restoring functional osteogenic differentiation.

Collectively, our findings position MSC-derived NGs as a new class of osteoimmunomodulatory nanotherapeutic that bridge immunoregulation and bone regeneration. Compared with cell-based therapies, NGs offer distinct advantages in safety, manufacturability, storage, and regulatory compliance. Unlike synthetic immunomodulatory biomaterials, NGs retain the complex, cell-derived molecular signature necessary to fine-tune immune responses in a context-dependent manner. Importantly, the ability to engineer NG function through parental cell conditioning provides a modular and programmable platform adaptable to diverse pathological microenvironments. Nevertheless, several limitations should be acknowledged. This study focuses on *in vitro* inflammatory models, and *in vivo* validation will be essential to assess biodistribution, persistence, and therapeutic efficacy within complex immune environments. Additionally, while IL-10 enrichment correlated with functional outcomes, NGs likely act through multiple synergistic pathways, including membrane-bound ligands and additional cytokines or microRNAs. Future research integrating multi-omics profiling and mechanistic studies will be necessary to fully elucidate NG’s mode of action.

## 5. Conclusion

In conclusion, our findings establish bioengineered MSC-derived NGs as a scalable and programmable acellular nanotherapeutic platform for osteoimmunomodulation. By combining high-yield NG fabrication with biomimetic microenvironment MSC conditioning, we demonstrate that hypoxic and three-dimensional cues synergistically encode immunoregulatory function into NGs, resulting in enhanced protection of MSCs under inflammatory stress, and partial restoration of osteogenic capacity under inflammatory conditions. These findings position NGs as active immunomodulatory agents capable of directing immune–bone crosstalk while overcoming key limitations of cell- and EV–based therapies, including scalability and translational feasibility. Collectively, this research introduces a modular strategy for engineering immunoregulatory nanotherapeutics and highlights NGs as a promising therapeutic avenue for the treatment of challenging inflammatory bone defects and skeletal pathologies.

## Acknowledgements

We would like to acknowledge Àlex Monte i Homs and Andrés Fernández Pilar for their assistance on this study.

## Notes

### Competing Interest Statement

The authors have declared no competing interest.

